# Mimetic butterfly wings through mimetic butterfly eyes

**DOI:** 10.1101/2023.10.05.561098

**Authors:** Andrew Dang, Gary D. Bernard, Furong Yuan, Aide Macias-Muñoz, Ryan I. Hill, J. P. Lawrence, Aline G. Rangel Olguin, Armando Luis-Martínez, Sean P. Mullen, Jorge Llorente-Bousquets, Adriana D. Briscoe

## Abstract

Color vision is thought to play a key role in the evolution of animal coloration, while achromatic vision is rarely considered as a mechanism for species recognition. Here we test the hypothesis that brightness vision rather than color vision helps *Adelpha fessonia* butterflies identify potential mates while their co-mimetic wing coloration is indiscriminable to avian predators. We examine the trichromatic visual system of *A. fessonia* and characterize its photoreceptors using RNA-seq, eyeshine, epi- microspectrophotometry and optophysiology. We model the discriminability of its wing color patches in relation to those of its co-mimic, *A. basiloides*, through *A. fessonia* and avian eyes. Visual modeling suggests that neither *A. fessonia* nor avian predators can readily distinguish the co-mimics’ coloration using chromatic or achromatic vision under natural conditions. These results suggest that mimetic colors are well-matched to visual systems to maintain mimicry, and that mate avoidance between these two look-alike species relies on other cues.

## Introduction

Müllerian mimicry is an effective defensive strategy against predators where multiple unpalatable species adopt the same conspicuous appearance and contribute to predators avoiding the conspicuous signal. Often accompanying these signals are chemical, physiological, and behavioral defenses. *Limenitis* butterflies provide several examples of such a phenomenon. For instance, *L. archippus* (viceroy) benefits from the chemical defenses of its co-mimics, *Danaus plexippus* (monarch) and *D. gilippus* (queen), as avian predators learn to associate distastefulness with the similar wing patterns of all three species^1,2^. Though mimicry can involve other sensory modalities such as the auditory mimicry observed in moths^3^, most studies of mimicry emphasize coloration and vision.

Studies of visual mimicry have generally focused on the targeting of prey by predators^4,5^. Examples include avian predators learning to avoid brightly colored frogs^6^ and aposematic butterflies^7–9^, and carnivores learning to avoid mimetic snakes^10^. However, the perceptual capabilities of prey and predator considered jointly have only been investigated relatively recently^11–13^. For most birds, moth and butterfly larvae and adults comprise a key component of their diet, and lepidopteran abundance is positively correlated with bird reproductive success^14^. Despite these studies, there is still a dearth in knowledge of the visual systems of lepidopteran prey and how these systems contribute to the evolution and maintenance of butterfly aposematism and mimicry.

This study focuses on the visual system of *Adelpha fessonia*, a member of a genus which consists of over 100 described species and a similar number (∼100) of subspecies^15^ ranging from the southern United States to South America. Recent phylogenies suggest that *Limenitis*^16^ and the Asian genus *Athyma*^17^ should be included in *Adelpha*. Among *Adelpha* adults, wing patterns involving multiple convergent coloration patterns suggest several mimicry complexes^15^. For example, at least seven *Adelpha* species in Venezuela have independently evolved an orange band bordering on and touching the white band of the dorsal hindwing^15^. A field study deploying paper and clay model butterflies and counting bird attacks found that mimetic *Adelpha* wing color patterns are an effective defense mechanism against predators^9^. This study and others^18,19^ have highlighted the frequency-dependent benefits conferred on the mimic by the model and have also revealed some nuances in the classification of an *Adelpha* species as either model or mimic.

Our focal taxon *Adelpha fessonia* has orange forewing tips with conspicuous white bands on both wings that likely act to reduce predation by signaling escape potential, unpalatability, and/or by disrupting prey recognition^20^. The white stripe may also influence mating success^21^. Like other *Adelpha* butterflies, *A. fessonia* fly in a quick and unpredictable manner, which may cause predators to expend more energy in catching the prey than they take in by obtaining them, thus associating butterflies with similar patterns as being too costly to bother catching^22–24^. *Adelpha fessonia* may be distasteful due to obtaining toxins as larvae from the plants they consume in the genus *Randia* (Rubiaceae)^15^. As adults, *Adelpha* may maintain unpalatability by feeding on white *Cordia* flowers, Boraginaceae plants that are hypothesized to contribute to the distastefulness of unpalatable species^25^. Across the *Adelpha* genus, larval host plant diversity is not only correlated with increasing *Adelpha* species diversity, suggesting an evolutionary arms race between *Adelpha* butterflies and their host plants, but also suggests that a potential plethora of toxic chemicals contributes to their unpalatability^16,26,27^.

Although many species of *Adelpha* share the general orange-tip and white band appearance, *A. fessonia* is unique in having the white stripe continuously unbroken from the costa to the anal margin^15^. A second species, *A. basiloides*, is similar in having some white expanding into the discal cell, with a small break in the white stripes differentiating the two. These species share common localities in the United States, Mexico^28^, Guatemala, Belize, El Salvador, Honduras, Nicaragua, Costa Rica, and Panama^15^ which, combined with their similar wing patterns, suggests they are mimics of each other. Their shared color pattern may lead to generalization from predators to avoid unprofitable prey. Strengthening the case for mimicry is that in Central America *A. basiloides’* white band is wide, matching *A. fessonia* and other taxa, but in western Ecuador, *A. basiloides’* white band is narrow, matching several other *Adelpha* species found there^15^.

For many butterflies in the family Nymphalidae, which includes species of *Adelpha/Limenitis*, vision is based on three opsin-based photoreceptors^29–32^. This number of photoreceptors, which underlies trichromatic color vision, is commonly found in flower foraging insects as seen in honeybees^33^, bumblebees^34^, and hawkmoths^29^. Typically, these photopigments are encoded by two short wavelength-sensitive opsin genes (*UVRh* and *BRh*) and one long wavelength-sensitive (*LWRh*) opsin gene^35,36^ although exceptions exist^37–39^. Opsin proteins together with a light-absorbing 11-*cis-3*- hydroxy retinal chromophore are the constituent parts of photoreceptor molecules that are sensitive to wavelengths of light in the UV-visible spectrum. Amino acid substitutions in the chromophore binding pocket domain of the opsin protein can spectrally tune the wavelength of peak absorbance (λ_max_) of the photopigment^40^. Additional photoreceptor classes can also be produced via opsin gene duplication or from single opsins expressed together with photostable filtering pigments that coat the rhabdom^41^.

Lepidoptera rely on vision to navigate and interact with their world. *Adelpha fessonia* is no exception. However, little is known about how the visual system of *Adelpha* butterflies affects their ability to detect potential mates whilst living in habitats shared by communities of co-mimics. A PCR-based survey of opsins expressed in adult eyes of *L. arthemis astyanax* and *L. archippus* yielded three opsin mRNA transcripts, encoding ultraviolet (UV)-sensitive *(UVRh)*, blue-sensitive (*BRh*) and long wavelength (LW)-sensitive (*LWRh)* opsin mRNAs, respectively^32^. Moreover, the potential for variation in opsins among these butterflies was demonstrated in a separate study where the λ_max_ values of the LW-sensitive rhodopsins were shown to differ by as much as 31 nm^31^.

In this study, we test the hypothesis that the butterfly *Adelpha fessonia* may be able to use achromatic cues or brightness vision to discriminate its wing colors from that of its co-mimic *A. basiloides,* while potential avian predators likely find these mimetic wing colors to be indiscriminable. To test this hypothesis, we characterize the visual system of *A. fessonia* and measure several parameters needed for visual modeling. Such a model of vision can help predict whether *A. fessonia* use coloration, brightness, or other cues such as pheromones to differentiate conspecifics from heterospecific mimics. We made RNA-seq libraries from eye and brain tissue of *A. fessonia* and related species and built transcriptome assemblies to identify their opsins. We then quantified opsin expression levels using RNA-seq data. We used epi- microspectrophotometry and optophysiology (a method which measures the decrease in intensity of eyeshine due to the movement of intracellular pigment granules in response to stimulating light) to measure the λ_max_ values of *A. fessonia* LW- and UV-sensitive photopigments, respectively. We inferred the λ_max_ value of the *A. fessonia* blue-sensitive photopigment using comparative sequence analysis together with functional expression data for *Limenitis* blue-absorbing rhodopsins. We then modeled the trichromatic color vision system of *A. fessonia* using the characterized photoreceptor spectral sensitivities. Color space modeling of the co-mimics’ wing coloration provides evidence that the coloration found on the orange patches and white stripes of *A. fessonia* and *A. basiloides* wings may be indistinguishable to both *A. fessonia* and avian predators under field conditions. Our efforts represent the first comprehensive study of the visual system of a butterfly species in the genus *Adelpha* and lay the foundation for future studies on the contribution of *Adelpha* butterflies’ vision to the evolution of anti-predator defenses including mimicry.

## Results

### *Adelpha fessonia* eyeshine is sexually monomophic

Although we did not examine eye size or visual acuity in *Adelpha fessonia*, traits that are often sexually dimorphic^42–44^, photographs of *A. fessonia* compound eyes and eyeshine do not show any obvious dimorphism between the two sexes (Fig. 1A-C). Eyes of both sexes are characterized by heterogeneity in the color of the eyeshine across the retina with blue, green, and yellow-green-reflecting ommatidia (Fig. 1E, F). This observation suggests their color vision system is sexually monomorphic. The lack of red-reflecting ommatidia, which are red in color because they contain a high density of blue-absorbing filtering pigments, also suggests an absence of red-shifted R3-9 photoreceptors cells in *A. fessonia*^45^. This is because many butterflies, including other nymphalid species such as *Heliconius*, have filtering pigments which coat the rhabdoms and filter short-wavelength light available to the long-wavelength sensitive rhodopsins, shifting the photoreceptor cell’s peak sensitivity towards the red part of the visible light spectrum^46,47^. In nymphalid butterflies without saturated red eyeshine, a previous study found R3-8 photoreceptor cells with peak sensitivities at or below 535 nm (green) light^48^. This does not preclude, however, a class of long wavelength-sensitive photoreceptor cell with a peak sensitivity >590 nm due to optical filtering of the basal R9 cell by the overlying R1-8 photoreceptor cells^45^. Even with the absence of eyeshine suggesting red-sensitive photoreceptor cells, it is still expected that *A. fessonia* will have UV-, blue- and LW-sensitive photoreceptor cells, corresponding to three kinds of opsin.

**Figure 1.**
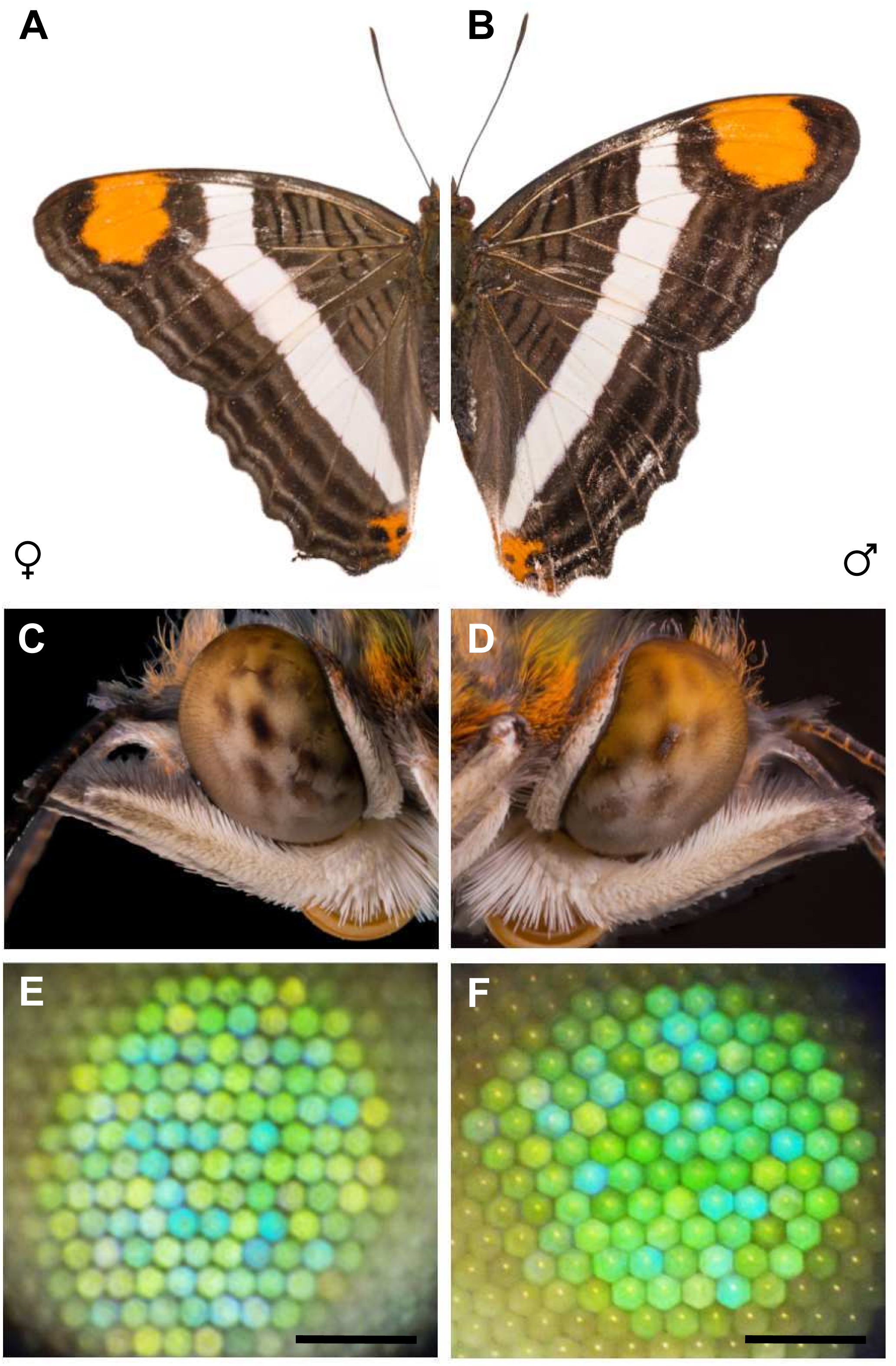
*Adelpha fessonia* wings, compound eyes, and eyeshine. **a** Photographs of an *Adelpha fessonia* female wing and **b** an *A. fessonia* male wing. **c** Photographs of an *A. fessonia* female eye and **d** an *A. fessonia* male eye. **e** Photographs of *A. fessonia* female eyeshine and **f** male eyeshine. Scale bar=100 μm.

### UV, blue and LW visual opsins of *Adelpha fessonia*

Indeed, our BLAST searches of the *A. fessonia* transcriptome assembly yielded three visual opsin transcripts corresponding to *UVRh, BRh*, and *LWRh,* and four additional transcripts encoding RGR-like opsin, unclassified opsin, pteropsin, and Rh7 opsin. A similar number of visual opsin transcripts (n=3) was found in the other surveyed *Adelpha/Limenitis* species except for *A. leucerioides,* where we found two LW opsin mRNA transcripts, *LWRh1* and *LWRh2,* and evidence of a recent gene duplication (Fig. 2C). Across all three visual opsin phylogenies, the same sister taxa were recovered, with the genus *Limenitis* recovered as a monophyletic clade with high bootstrap support (Fig. 2). A site-specific maximum likelihood model showed no evidence of positive selection among the opsin genes of *Adelpha*.

**Figure 2.**
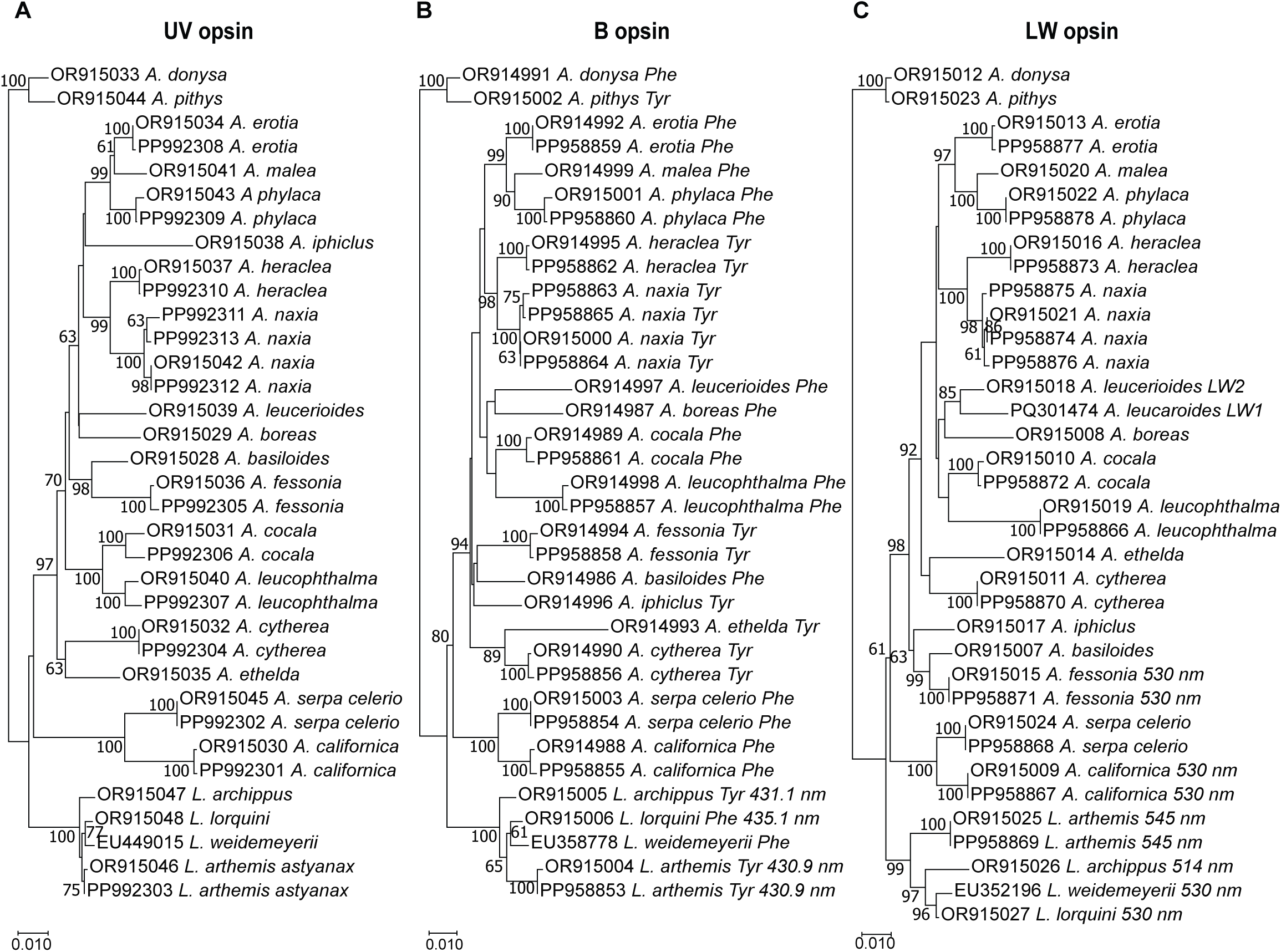
Phylogenies of *Adelpha* UV, blue, and LW opsin-encoding mRNA transcripts and wavelength of peak absorbance (λ_max_) of the encoded blue- and LW-absorbing rhodopsins. **a** *UVRh*, **b** *BRh*, and **c** *LWRh* maximum likelihood nucleotide phylogenies reconstructed using full-length coding sequences, the HKY85 model, and 500 bootstrap replicates. Bootstrap support values >60% are shown. **b** Blue opsin amino acid at site 195, either phenylalanine (Phe) or tyrosine (Tyr), and λ_max_ values of the blue-absorbing rhodopsins, where known^49^. **c** LW-absorbing rhodopsin λ_max_ values, where known^30^. All opsin transcripts appear to be single-copy except for *A. leucerioides LWRh,* which is duplicated. Specimen locality information for the sampled butterflies is given in Supplementary Data 1. Scale bar = substitutions/site.

### Visual opsin mRNAs are highly expressed in *A. fessonia* heads

We found that the *UVRh*, *BRh* and *LWRh* opsin mRNAs were highly expressed in *A. fessonia* head tissue while RGR-like opsin, pteropsin, and RH7 were not (Fig. 3). *LWRh* opsin mRNA had the highest expression followed by the *BRh* and *UVRh* opsin mRNAs with similar expression levels. This is to be expected as these three opsin mRNAs are expressed in the compound eyes of *Limenitis* ^49^. The unclassified opsin, a candidate retinochrome, was intermediate in expression compared to the visual opsins and pteropsin, RGR-like opsin, and Rh7.

**Figure 3.**
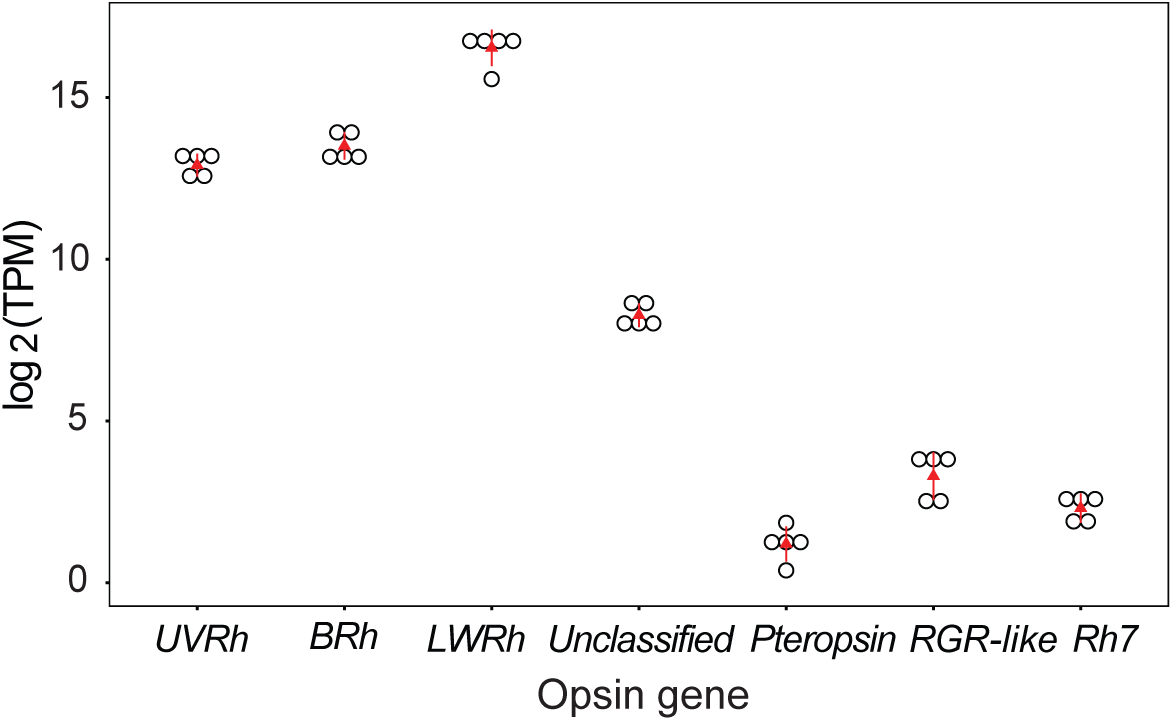
Expression levels of *Adelpha fessonia* opsin genes in adult head (eye + brain) tissue. mRNA transcripts measured using RNA-seq data (n=5 individual butterflies) and kallisto quantification. Y-axis is in transcripts per million on a log_2_ scale. Triangles indicate median values and open circles indicate individual data points.

### λ_max_ values of *Adelpha fessonia* LW and UV rhodopsins

*Adelpha fessonia* and *A. californica* LW rhodopsins from dark adapted and partially bleached eyes as measured using epi-microspectrophotometry show peak absorbances at 530 nm (Fig. 4A) similar to the λ_max_ values of *Limenitis lorquini* and *L. weidmeyerii*^30,31,49^.

**Figure 4.**
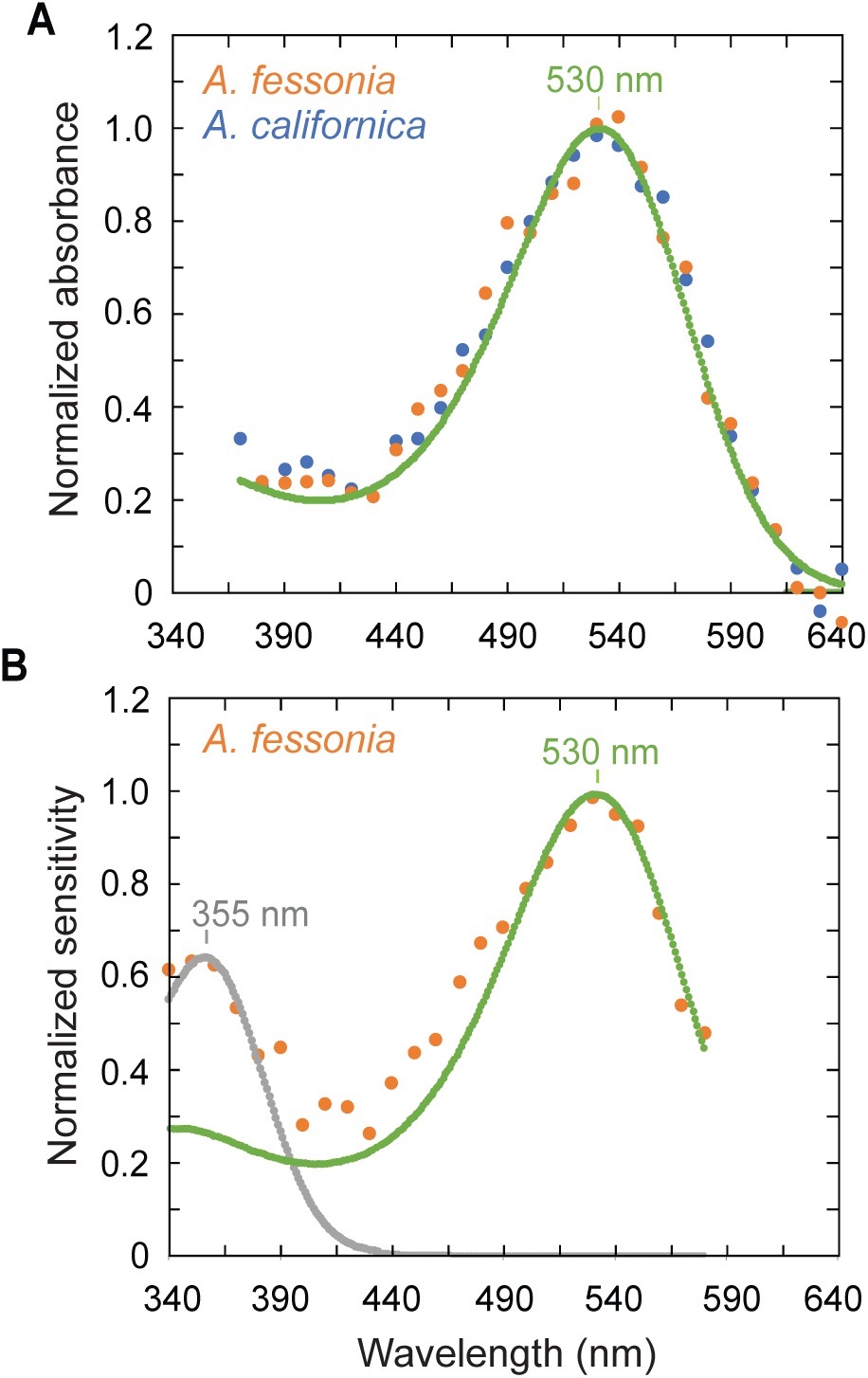
Epi-microspectrophotometry and optophysiology of *Adelpha fessonia* eyes. **a** Normalized absorbance spectra of *Adelpha fessonia* and *A. californica* long wavelength-absorbing rhodopsins measured by epi-microspectrophotometry. Experimental spectra of *A. fessonia* (n=1 butterfly)(orange dots) and *A. californica* (n=1 butterfly)(blue dots). LW rhodopsin template with peak absorbance at 530 nm (green line). **b** Normalized pupillary sensitivity of *A. fessonia* (n=1 butterfly)(orange dots). The contributions of UV- (grey dotted line) and LW- (green dotted line) sensitive rhodopsins to the pupillary sensitivity are indicated by fits to a computational model which includes both UV (λ_max_=355 nm) and LW (λ_max_=530 nm) rhodopsins.

### Pupillary sensitivities of *A. fessonia* in the UV and long wavelengths

The pupillary sensitivity data of adult *A. fessonia* eyes allowed us to independently estimate the wavelength of peak absorbance of the UV and LW rhodopsins (Fig. 4B). The good fit of a 355 nm rhodopsin template to the pupillary sensitivity data shows that only the UV-sensitive 355 nm photoreceptors contribute to responses in the range 340-380 nm. Similarly, the fit of a 530 rhodopsin template from 490-580 nm shows that only the green-sensitive 530 nm receptors contribute to long-wavelength sensitivity, confirming the epi-microspectrophotometry estimate that the peak absorbance of the LW rhodopsin of *A. fessonia* is 530 nm. In the range 390-480 nm the elevated sensitivity of data argues for a minor contribution from a blue-sensitive photoreceptor, but its spectral peak could not be determined from these experiments. Pigments in the 530 nm region are the most abundant and easiest pigments to measure due to their expression in R3-8 photoreceptor cells^49^, hence the higher sensitivity when compared to the peak near R355. The experimentally determined 355 nm and 530 nm peaks of the *A. fessonia* eye’s physiological response to light are driven by the UV- and LW-sensitive photopigments. The differing abundances of each pigment and the photoconversions of each rhodopsin to metarhodopsin (which has a peak in the bluish range) make it difficult to measure the λ_max_ value of the blue-absorbing rhodopsin.

### The blue-absorbing rhodopsins of *Adelpha/Limenitis*

To bypass technical challenges of measuring the blue rhodopsin *in vivo* using either epi-microspectrophotometry or optophysiology, we aligned and compared the blue opsins of 22 species in the *Limenitis/Adelpha* genera. Then we used absorbance spectra data from HEK293-expressed rhodopsins of species in the *Limenitis* clade to infer the absorbance peak of the *Adelpha fessonia* blue-absorbing rhodopsin, including data from a previously unstudied species, *L. lorquini,* a close relative of *A. fessonia*.

The blue-absorbing rhodopsin of *Limenitis lorquini* has a peak absorbance at 435.1 nm and a phenylalanine (F) at the known spectral tuning site, amino acid 195 (Fig. 5A,D). *Limenitis arthemis* and *L. archippus* blue-absorbing rhodopsins have peak absorbances of 430.9 nm and 431.1 nm, respectively, and both have a tyrosine at amino acid 195 (Fig. 5B and D)^49^. A substitution of tyrosine to phenylalanine at amino acid 195 (Y195F) in the *L. arthemis* blue opsin results in a red-shifted photopigment with a λ_max_ of 435.1 nm (Fig. 5C). As seen from the amino acid alignment in Fig. 5D, *A. fessonia* has tyrosine at position 195 suggesting that the blue-absorbing rhodopsin of *A. fessonia* has a peak absorbance at around 431 nm. Although it is possible that the *A. fessonia* blue-absorbing rhodopsin has a different wavelength of peak absorbance than suggested by this comparative study, this is unlikely due to the high sequence similarity (97.3% identity or 321 out of 330 amino acids) of the *A. fessonia* blue opsin compared the *Limenitis* blue opsins that have been functionally expressed.

**Figure 5.**
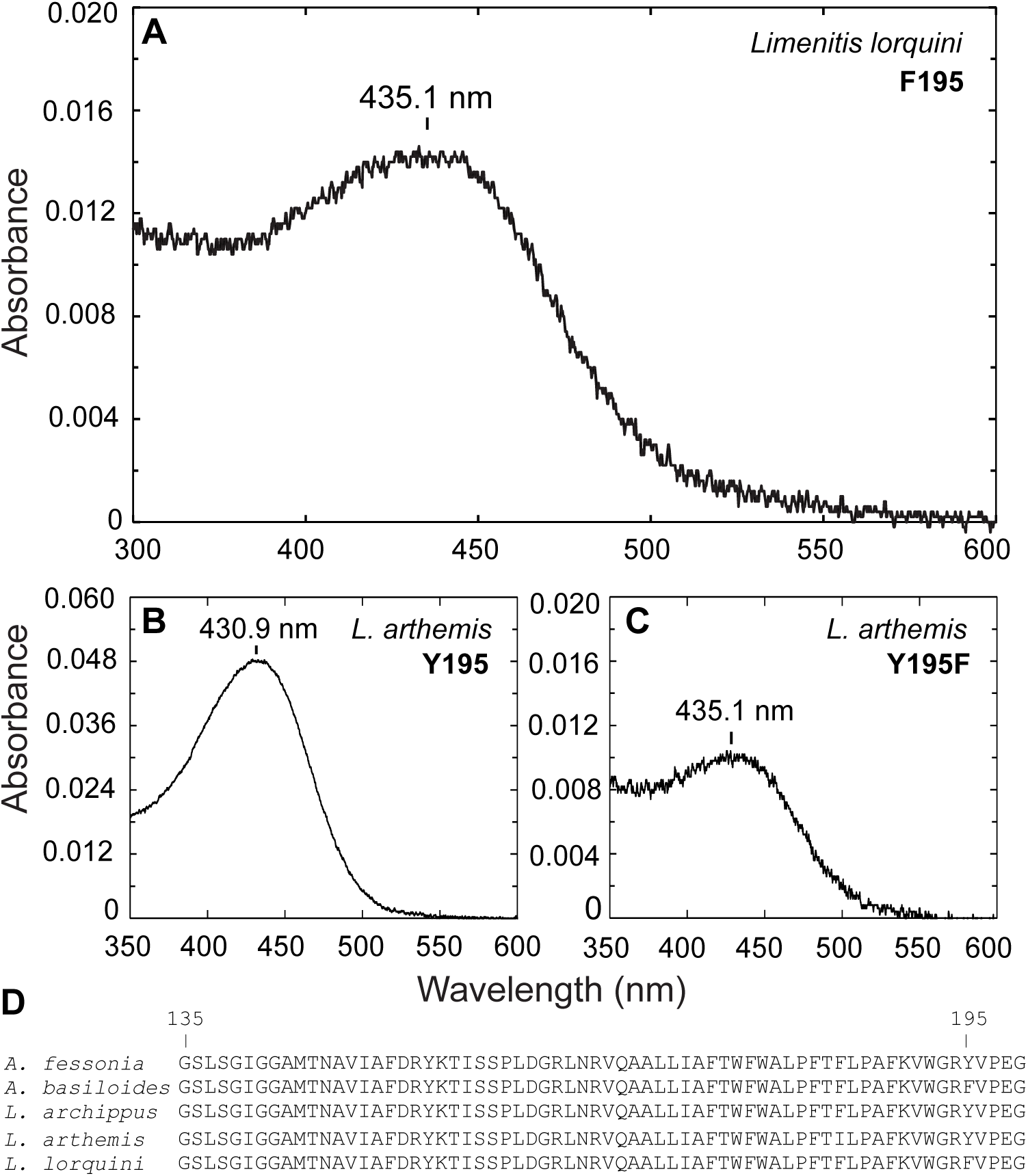
*Limenitis* blue-absorbing rhodopsin dark spectra based on heterologous expression in HEK 293 cells and known spectral tuning sites. **a** Dark spectra of wildtype *Limenitis lorquini* (n=1 independent experiment), **b** wildtype *L. arthemis* (n=1) and **c** Y195F mutant *L. arthemis* (n=1) blue-absorbing rhodopsins. Spectra of **b** and **c** reproduced with permission from Frentiu et al. (2015)^49^. Each construct was expressed 2-4 times in independent experiments with 11-*cis*-retinal, purified, and measured, and the experiment with the highest optical density was selected for further analysis. Each panel represents the average of 5-7 measurements (technical replicates) from the single best independent experiment for that construct. **d** Partial amino acid alignment of the blue opsin of representative species indicating the location of two amino acid sites, 135 and 195, with known spectral tuning effects^49,96,97^.

### *Adelpha fessonia* and avian predators may not distinguish mimetic *Adelpha* wing colors

The dorsal white stripes and orange patches of most *Adelpha* wings look highly similar to the human eye between species (Fig. 6A) and so we investigated if this is also true for *A. fessonia* and its avian predators. First, we examined whether wing reflectance is sexually dimorphic in *A. fessonia* or *A. basiloides* and found little evidence for this (Supplementary Figure 1), so we pooled male and female measurements in subsequent analyses. The white stripes on the dorsal wings of *A. basiloides* have similar reflectance in the 300 to 700 nm range compared to *A. fessonia* wings while the orange patches overlap from 300-600 nm and then diverge from 600-700 nm (Fig. 6B).

**Figure 6.**
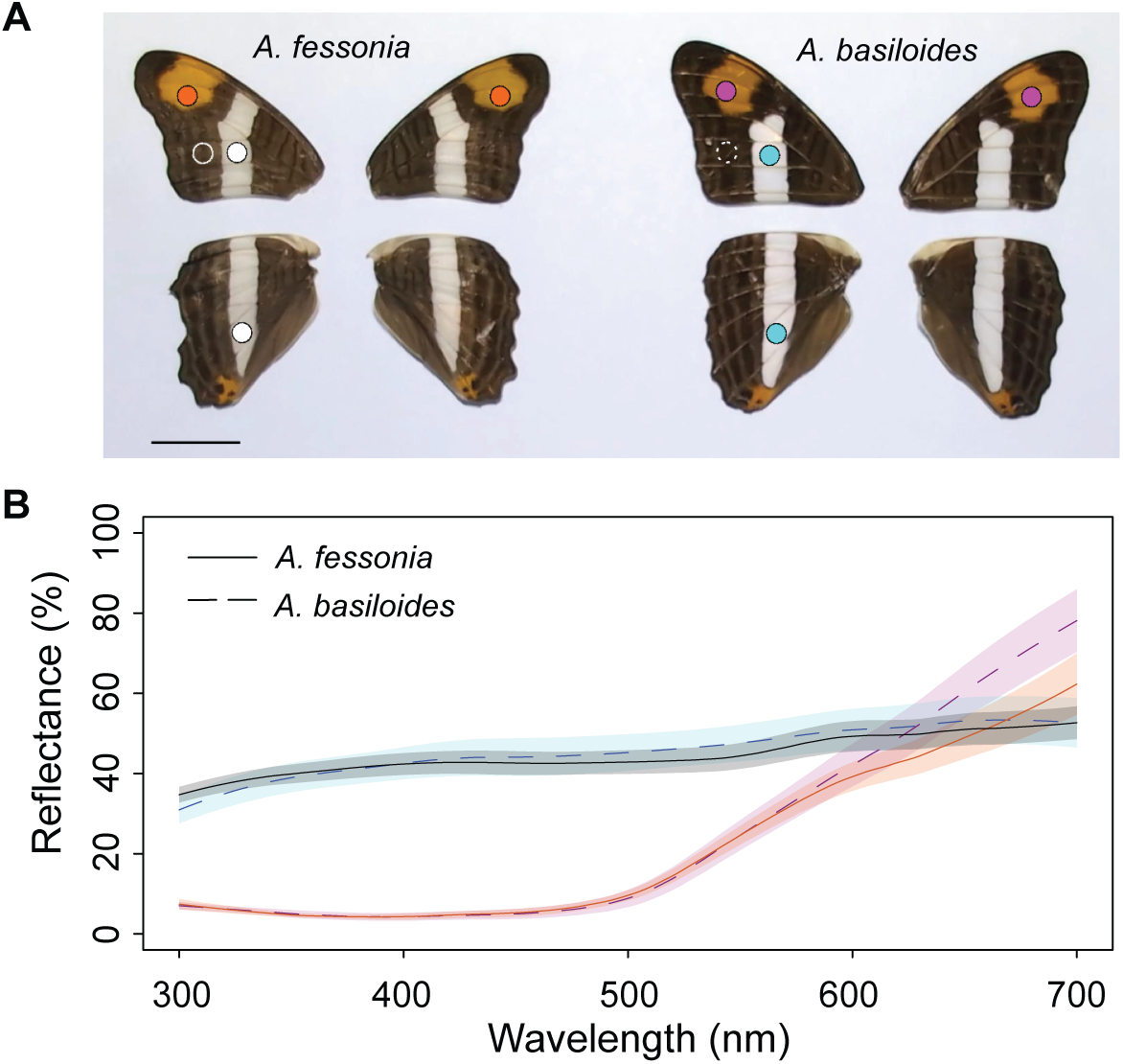
Photographs and wing reflectance spectra of *Adelpha fessonia* and *A. basiloides* mimetic butterflies. **a** Photographs of dorsal wings of *A. fessonia* (left) and *A. basiloides* (right) with scale bar representing 1 centimeter. Circles indicate parts of the wings where reflectance spectra were measured. Orange dots=*A. fessonia* left and right dorsal orange forewing patches; white dots=*A. fessonia* left dorsal forewing and hindwing white stripes; white circle, solid line=*A. fessonia* left dorsal forewing brown; pink dots=*A. basiloides* dorsal left and right orange forewing patches; blue dots=*A. basiloides* left dorsal forewing and hindwing white patches; white circle, dashed line=*A. basiloides* left dorsal forewing brown. **b** Reflectance measurements taken from *A. fessonia* left (n=13 independent measurements) and right (n=12) dorsal orange forewing patches (orange dots in **a**), *A. fessonia* left dorsal forewing (n=13) and hindwing (n=10) white stripes (white dots), *A. basiloides* left (n=18) and right (n=17) dorsal orange forewing patches (pink dots), and *A. basiloides* left dorsal forewing (n=13) and hindwing (n=14) white wing stripes (blue dots). Shaded areas represent standard deviations and lines (solid, *A. fessonia*; dashed, *A. basiloides*) represent means.

The orange and white dorsal wing colors of *A. fessonia* and *A. basiloides* form clusters in the color spaces of the butterfly and the bird visual systems, suggesting these colors are very similar in the perceptual spaces of both the butterfly and its potential predators (Fig. 7 A-C). For none of the three achromatic visual models did the mean discriminability (JND) between the orange patches or between the white stripes of the two butterflies exceed a threshold of one JND (Fig. 7 D-F, right). This is also the case for two of the three chromatic visual models comparing the butterflies’ oranges and forewing dorsal whites, respectively (Fig. 7D-F, left). The only exception was the violet-sensitive bird visual model, in which mean JNDs for the forewing oranges exceeded one (Fig. 7E, left). A discriminability threshold of one assumes idealized conditions such as those found in the lab rather than those found in nature. A bird with a violet-sensitive visual system of the kind we model here might be able to discriminate these oranges in a laboratory setting but may be unable or indifferent to these color differences in the wild. Overall, these results suggest that *A. fessonia* and *A. basiloides* are close mimics in the color spaces of both butterflies and birds. As predictive tools, these models will require behavioral studies to confirm our conclusions as well as further refinements in the parameter estimates used in the visual models (see below).

**Figure 7.**
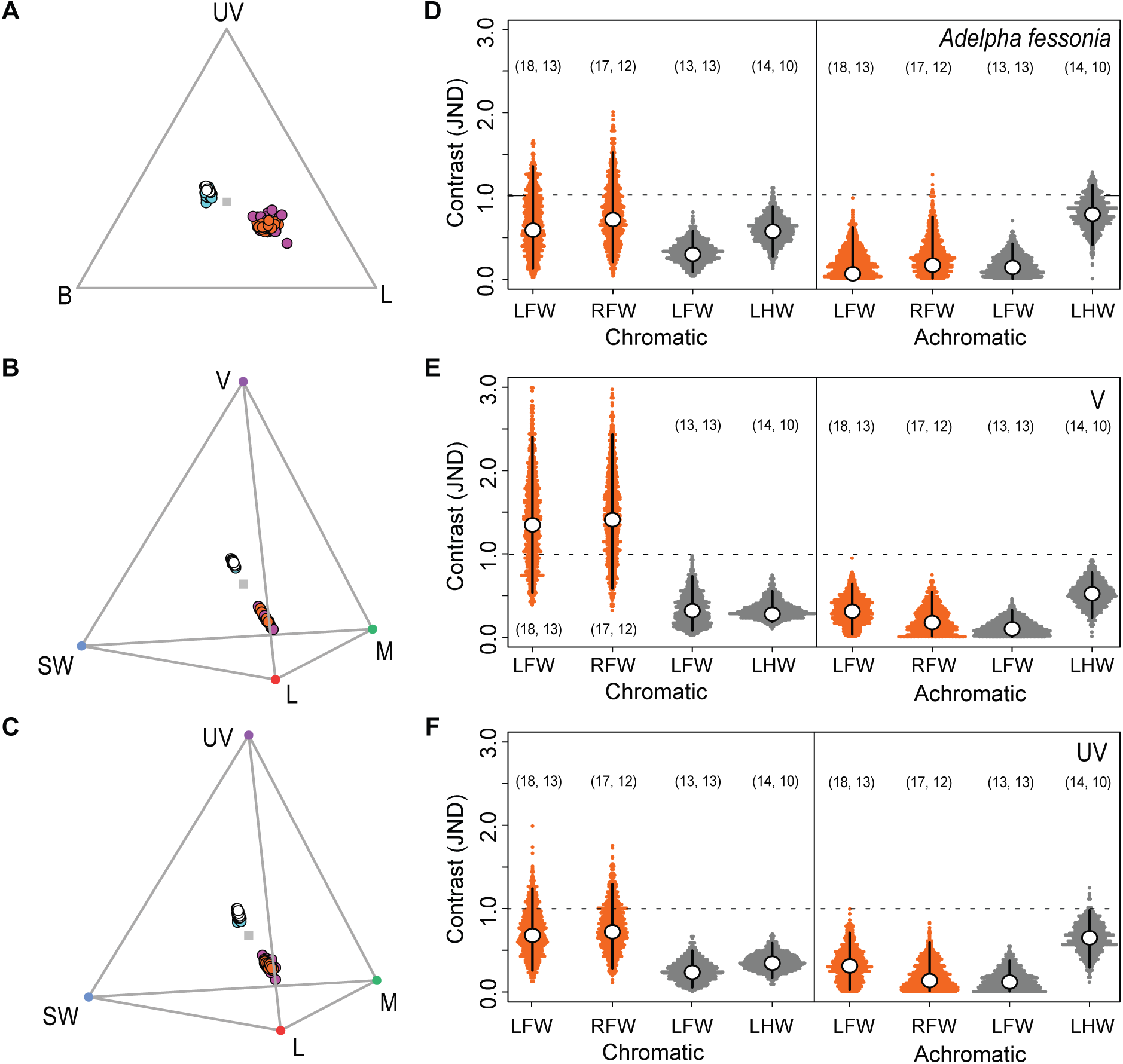
Wing reflectance spectra of *Adelpha fessonia* and *A. basiloides* plotted in the trichromatic color space of *Adelpha fessonia* and the tetrachromatic color spaces of the UV-sensitive and violet-sensitive bird visual systems, and just noticeable differences (JNDs) estimated from the corresponding visual models. The dorsal orange patches and white stripes of mimetic *Adelpha fessonia* and *A. basiloides* butterflies are likely indistinguishable to *Adelpha fessonia* and their avian predators in the wild. The dorsal orange patch of *A. fessonia* (orange dots, n=25 independent measurements) and *A. basiloides* (pink dots, n=35), and the dorsal white stripes of *A. fessonia* (white dots, n=23) and *A. basiloides* (blue dots, n=27) mapped in the trichromatic color space of **a** the *A. fessonia* visual system, and the tetrahedral color spaces of **b** the violet-sensitive and **c** the UV-sensitive, avian visual systems. For *A. fessonia,* UV=ultraviolet, B=blue and L=LW rhodopsin. For birds, UV=ultraviolet, V=violet, SW=blue, M=RH2, L=LWS rhodopsin. Right: Distances between the two species’ orange (orange dots) and white (white dots) wing color patches as viewed through **d** the *A. fessonia* visual system, **e** the violet-sensitive, and **f** the UV-sensitive avian visual systems. Color distances are in units of chromatic contrast (JND) between *A. fessonia* and *A. basiloides* wing colors with a JND threshold (dotted line) of 1. Open circles represent mean JND values estimated from 1000 bootstrapped replicates with black bars representing 95% confidence intervals. Orange represents individual bootstrapped JNDs derived from comparisons of *A. fessonia* and *A. basiloides* orange wing colors and dark grey represents individual bootstrapped JNDs derived from comparisons of *A. fessonia* and *A. basiloides* white wing colors. Sample sizes of wing reflectance spectra used to estimate these mean JNDs via bootstrapping are indicated in parentheses with numbers for *A. fessonia* on the left and numbers for *A. basiloides* on the right. LFW=left forewing, RFW=right forewing, LHW=left hindwing.

## Discussion

Our experiments have allowed us to explore the hypotheses that *Adelpha fessonia* is able to discriminate its wing colors achromatically from that of its co-mimic *A. basiloides,* while birds are unable to discriminate the co-mimics’ colors using either chromatic or achromatic cues. Studies in color vision typically focus on the impact of chromatic cues and control for achromatic information (Kelber et al. 2003). However, most visual stimuli have a combination of chromatic and achromatic cues, and both should be accounted for when making predictions (Olsson et al. 2018). *Papilio xuthus* foraging behavior, for instance, has been shown to make use of achromatic cues to elicit behavioral responses^50^.

To explore these hypotheses, we needed to conduct physiological experiments which would allow us to measure or estimate several parameters of the *A. fessonia* visual model: number of visual opsins, wavelength of peak sensitivity of its R1-8 photoreceptor cells, relative ratio of each spectral class of photoreceptor, and wing reflectances of the co-mimics (Table 1, Fig. 6 and Supplementary Figure 1). We began by characterizing the photopigments of *Adelpha fessonia* eyes. We showed that *A. fessonia* have three kinds of opsins and the wavelengths of peak sensitivity for each of these photopigments is consistent with a visual system with the potential for trichromacy. We showed that *A. fessonia* eyeshine (and likely color perception) is not obviously sexually dimorphic, although we did not quantify eye size or visual acuity, traits which often are sexually dimorphic. We found that *A. fessonia* is unlikely to have color vision in the red range (600-700 nm) due to a 535 nm LW rhodopsin with a low absorbance above 600 nm (Fig. 4A) and an absence of red-reflecting ommatidia. Taken together, these results suggest the visual system of *A. fessonia* is indeed trichromatic.

**Table 1.** Parameters used in the visual models and their estimates from anatomical, physiological or behavioral data.

| Visual model |  |  |  |
| --- | --- | --- | --- |
| Peak sensitivities (nm) | *Adelpha fessonia*<br>355, 431, 530<br>(UV, S, L) | UV bird<br>372, 449, 502, 563<br>(UV, S, M, L) | Violet bird<br>418, 453, 507, 571<br>(V, S, M, L) |
| References | This study | Hart et al. 2000<sup>98</sup><br>(blue tit) | Bowmaker et al. 1997;<br>Carvalho et al. 2007<br>(chicken)<sup>99,100</sup> |
| Method | Epi-microspectrophotometry;<br>optophysiology;<br>heterologous<br>opsin expression, UV-visible spectroscopy &<br>comparative sequence<br>analysis | Microspectrophotometry | Microspectrophotometry;<br>Heterologous opsin<br>expression & UV-visible<br>spectroscopy |
| Photoreceptor density | 0.15:0.13:1 (uv:s:l) | 1:2:2:4 (uv:s:m:l) | 1:2:4:4 (v:s:m:l) |
| References | Frentiu et al. 2015<sup>49</sup>;<br>This study<br>(*Limenitis arthemis*) | Maier & Bowmaker<br>1993 (*Leiothrix lutea*);<br>Vorobyev & Osorio<br>1998<sup>78,81*</sup> | Kram et al. 2010<sup>86</sup><br>(chicken) |
| Method | Opsin *in situ*<br>hybridization | Microspectrophotometry | Brightfield & fluorescent<br>microscopy |
| $\omega$ (chromatic) | 0.13, 0.06, 0.12 | 0.10 | 0.06 |
| References | Hempel de Ibarra et al.<br>2001; Vorobyev et al.<br>2001 (honeybee)<sup>90,91</sup> | Maier 1992<br>(*Leiothrix lutea*)<sup>82</sup><br>Lind et al. 2013a<br>(*Melopsittacus*<br>*undulatus*)<sup>84</sup><br>Silvasti et al. 2021<br>(blue tit)<sup>83</sup> | Olsson et al. 2015, 2018<br>(chicken)<sup>87,88</sup> |

| Method | Electrophysiologyd<br>(Spectral sensitivity) | Behavior<br>(Spectral sensitivity) | Behavior<br>(Object discrimination) |
| --- | --- | --- | --- |
| $\omega$ (achromatic) | 0.16 | 0.18 | 0.22** |
| References | Srinivasan and Lehrer<br>1988 (honeybee)<sup>92</sup> | Lind et al. 2013b<br>(*Melopsittacus*<br>*undulatus*)<sup>85</sup> | Hodos et al. 1985;<br>Lind et al. 2013b<br>(*Columba livia*)<sup>85,89</sup> |
| Method | Behavior<br>(Contrast sensitivity) | Behavior<br>(Brightness<br>discrimination) | Behavior<br>(Brightness<br>discrimination) |

\* Vorobyev & Osorio 1998 appears to be the first paper to estimate these relative abundances of photoreceptor types for a bird with an uv-type visual system<sup>78</sup>. However, the source of data for this estimate is Table 1 in Maier & Bowmaker 1993<sup>81</sup>, which reports how many cells with a particular spectral sensitivity they were able to measure. This is an indirect measure of relative receptor abundance. It would be preferable to use an anatomical method in which all cell types in a photomicrograph of retina were scored based on oil droplets, opsin *in situ* hybridization or immunohistochemistry.
\*See Olsson et al. (2018) for estimates (0.20, 0.25 and 0.40) from other studies<sup>88</sup>.

We quantified opsin gene expression levels in *A. fessonia* and counted photoreceptor abundances from a close relative, *Limenitis arthemis*. Characterizing the UV and LW photopigments of *A. fessonia* eyes (and the blue-absorbing rhodopsin of several *Limenitis* species) allowed us to build a model of *A. fessonia* vision and provide us with hints of their perceptual world. While most of these experiments were performed on *A. fessonia*, for some parameters (relative abundance of each photoreceptor class, λ_max_ of the blue absorbing rhodopsin), it was necessary to make estimates from *Limenitis* butterflies due to difficulties we had in obtaining live pupae. Other estimates, such as measuring achromatic and chromatic weber fractions for *A. fessonia*, were similarly outside the scope of the study due to difficulty in obtaining live pupae among other challenges (i.e. our previous attempts to elicit visual feeding behavior from *Limenitis* butterflies in the lab did not work). Nonetheless, our findings are relevant to studying the evolution and ecology of visual perception in *Adelpha* butterflies more broadly.

The presence of heterogeneous eyeshine in butterfly eyes consisting of yellow- and red-reflecting ommatidia is not universal^48,51^. While eyeshine could be locally uniform, there are instances where heterogeneity occurs that could be indicative of adaptations that optimize spectral discrimination in the yellow, orange, and red range of the visible light spectrum. The patterns resulting from this heterogeneity can vary greatly between species though some species may share similar motifs. Lycaenidae, and certain nymphalid subfamilies, Nymphalinae and Charaxinae, share a red and yellow motif with the nymphalid subfamily Heliconiinae^52^. This red eyeshine in *Heliconius* butterflies is indicative of the presence of blue-absorbing “red” filtering pigments which coat the rhabdomeres in the R5-8 cells, and which cause a narrowing and red-shifting in the sensitivity of a photoreceptor cell expressing a green-absorbing rhodopsin^45,46^. The presence of, and heterogeneous expression of these red filtering pigments have been demonstrated to correlate with behavioral differences between butterfly species that have them and those that do not. *Heliconius erato,* a species possessing red filtering pigments, was able to differentiate between red (620 nm) and orange (590 nm) stimuli. *Vanessa atalanta,* a species without red filtering pigments, and with a 530 nm green receptor similar to that of *Adelpha fessonia,* was not^29,46^. *Adelpha fessonia* has a yellow/green/blue eyeshine pattern like what is seen in other nymphalids such as *Hypolimnas bolina,* denoting a lack of a red photoreceptor^51^.

It was expected that *A. fessonia* vision would have three major spectral classes of photoreceptors as described in other nymphalids^35^. We also expect genes involved in vision to be highly expressed in head tissue. RNA-seq quantification showed high expression levels of the three light-sensitive opsin mRNAs (*LWRh*, *BRh*, and *UVRh*) that are directly involved in vision along with an unclassified opsin that is also expressed in nymphalid butterfly secondary pigment cells (Fig. 3)^53^. The *A. fessonia* eye and brain has a relatively low level of pteropsin, RGR-like opsin, and RH7 mRNA expression by comparison.

We found the peak absorbances (λ_max_) of *A. fessonia* photopigments in the ultraviolet and long wavelength portions of the light spectrum corresponding to 355 nm and 530 nm, respectively, based on optophysiology and epi-microspectrophotometry of living eyes. For the blue portion of the light spectrum, we identified a photopigment with a peak sensitivity likely corresponding to 431 nm based on comparative sequence analysis (Figs. 4 and 5). Previously, amino acid substitutions at two sites were shown to be crucial for spectral tuning in blue-absorbing visual pigments in the small white butterfly, *Pieris rapae*^40^. In the case of *P. rapae*’s blue-absorbing pigment, a F to Y at position 177 caused a 4 nm blue shift. Conversely, site-directed mutagenesis of the opposite mutation (Y to F at site 195) in *Limenitis* BRh opsins caused the same magnitude of effect, although in the opposite direction (Fig. 3C-E and Frentiu et al. 2015^49^). Importantly, functional expression of wildtype *L. lorquini* and mutant *L. arthemis astyanax* opsins, which both have a F at site 195 have the same peak absorbance of 435 nm, while wildtype *L. arthemis astyanax* and *L. archippus,* with a Y at site 195 both have peak absorbances at 431 nm. Given that *A. fessonia* BRh shares a Y at site 195 with *L. arthemis* and *L. archippus*, it is highly likely its blue photopigment has a spectral peak at 431 nm, while that of *A. basiloides,* having an F at this site is likely to have a peak closer to 435 nm. The true λ_max_ values of the *A. fessonia* and *A. basiloides* blue-absorbing rhodopsins, which differ from *Limenitis* opsins at 8 additional amino acid sites out of 330, remain to be directly measured.

Do these small spectral tuning effects have an impact on these butterflies’ fitness? If so, then it is likely to be very small as we did not find any evidence of positive selection occurring in the blue opsins of the *Adelpha* lineage via a PAML site-specific test. The evolutionary reason behind why some species have tyrosine at site 195 and others have a phenylalanine is currently unknown but raises the question of whether or not this small tuning effect has an impact on the butterflies’ visual capabilities. What are the evolutionary benefits of one phenotype versus another and why do some members of the *Adelpha*/*Limenitis* clade possess the tyrosine or the phenylalanine phenotype? How does their vision affect their color pattern and host plant diversity? Although differing peak sensitivities in *Adelpha* BRh photoreceptors could cause variation in their visual perceptions which might play a role in creating and maintaining the color of objects they interact with, available evidence suggest that BRh is evolving neutrally across *Adelpha/Limenitis* species.

Butterflies are visually dependent creatures capable of discerning a plethora of colorful light stimuli^54,55^. Behaviors such as foraging for nectar and locating host/food plants have not been studied in *A. fessonia,* but with these data, the groundwork has been set to examine the role that chromatic and achromatic cues play in this system. Our results could also potentially lead to better understanding of how mimetic nymphalids are able to correctly identify conspecifics for mating by highlighting the need for future studies into the chemical cues that differ among species.

*Adelpha fessonia* eyeshine and wing color patterns (in the visible spectrum) are not apparently sexually dimorphic (Fig. 1A-F). Due to *A. fessonia’s* lack of a red-shifted green-absorbing rhodopsin and a blue-absorbing filtering pigment, its visual system resembles the UV-blue-green trichromatic visual system of honeybees, which are unable to discriminate monochromatic light above 550 nm^56,57^. Our JND results further suggest that *A. fessonia* may not be able to distinguish the orange patches on their own wings versus those on another similar-looking species like *A. basiloides*. Even if their tiny R9 cell is red-shifted due to optical filtering by overlying photoreceptor cells, the orange wing reflectance spectra between the two species are likely to be too similar to be distinguishable to *A. fessonia* because their 530 nm rhodopsin has little sensitivity above 620 nm, where the butterflies’ orange reflectance spectra begin to diverge (Fig. 4A and 6B) — a prediction that will need to be confirmed by behavioral experiments. Under either scenario, the orange aposematic signal is most likely used to ward off predators^24^.

Our modeling suggests that *A. fessonia* and UV-sensitive avian predators should not be able to discriminate between the orange patches or white stripes of *A. fessonia* and *A. basiloides,* chromatically or achromatically. Violet-sensitive avian predators might able to discriminate between the two species’ orange forewing colors under the idealized, bright light conditions of the laboratory but because the mean JNDs for these colors are still below 1.5, they may find this task difficult in the visually complex background of a forest where these butterflies live.

There is a vested interest for butterflies to distinguish conspecifics from co-mimics. Quickly finding conspecifics can improve mating success by lowering the time or energy needed for courtship. Studies of male Drosophila, for example, have shown that indiscriminate courtship results in lower lifetime mating success compared to either selective courtship or learning in the context of courtship^58^. In addition, selective courtship might also result in higher fitness as hybridization between species can result in a lower fitness for hybrid offspring such as the reduced feeding abilities observed in hybrid *Limenitis* butterflies^59^.

In this study, we have characterized the eyeshine and photopigments of *Adelpha fessonia* and modeled their visual capabilities as well as the visual capabilities of potential avian predators. Using the measured peak sensitivities of *A. fessonia* photopigments, we created a trichromatic color space model to quantify their potential color discrimination abilities. Limitations of the study include a lack of experimentally determined chromatic and achromatic weber fractions for *Adelpha fessonia* (or any other butterfly) and a similar lack of experimentally determined parameters from the same species of bird representing the UV- and violet-sensitive bird visual systems (see Methods and Table 1). Photoreceptor counts from *A. fessonia* and recordings from their blue-sensitive photoreceptor cells are also needed to verify our preliminary estimates reported here. Parameters used in visual models can have a big impact on results. For this reason, extra care should be made to ensure that parameters are based on either physiological or behavioral experiments where available, especially in comparisons across visual systems. Even so, the physiological data for the *A. fessonia* visual system we have presented here is a useful starting point for investigating those stimuli in the natural environment to which these butterflies may respond. Finally, the behavioral and ecological significance of this physiological data needs further investigation. Even though modeling is a powerful predictive tool, behavioral assays are still needed to confirm *A. fessonia* and bird color-discerning capabilities. Comparison of these species’ eyes with those of other butterflies can help to expand upon our knowledge of how a prey’s sensory system contributes to the generation and maintenance of biodiversity.

## Materials and Methods

### Photographs of butterflies

Whole mount specimen photos were taken with a Canon 5D Mk III and Canon 100mm f/2.8L macro lens with an off camera diffused flash. Each head photo is a stack of 50 photos taken with a Canon 5D Mk III and Canon MP-E 65mm 1-5x macro lens under constant, diffused lighting. Plane of focus was incrementally moved from the front to the back of the head with an automated slider rail. Photos were stacked with Zerene Stacker v.1.04 (Zerene Systems LLC, Richland). Final composite photos as well as whole mount photos were edited with Adobe Photoshop Lightroom 6 to adjust exposure and correct color.

### Photographs of butterfly eyeshine

The apparatus for eyeshine photos was the same as for epi- microspectrophotometry^48^. Briefly, it is a Leitz Ortholux-Pol microscope equipped with a Leitz 620-257 low stray-light Epi-illuminator, fed by a 45 w quartz-halogen lamp in an Ealing housing. Optics were a Leitz 8X/0.18P objective and 16X eyepieces. A Uniblitz shutter controlled exposure time. An Olympus TG-1 digital camera was attached to an eyepiece. Settings of the camera were: macro; iso 1600; zoom 4.0; focal length 18; metering mode 3; F/4.9; exposure program 2; color profile sRGB IEC61966-2.1; and 1920 x 2560 pixels. An intact butterfly (n=1 female and n=1 male) was mounted in a slotted plastic tube fixed to the goniometric stage, then oriented to set the eye’s direction of view. The microscope was adjusted to center the eyeshine patch in the field of view, focused on the cornea. After several minutes of dark-adaptation, a photo was taken using a 0.5 sec flash. The female eyeshine photograph was taken at 16:55 and the male eyeshine photograph was taken at 17:25. No eyeshine heterogeneity was observed across the retina.

### Specimen collection, RNA extraction, Sanger sequencing and RNA-seq

Ebel et al. (2015)^16^ demonstrated that Eurasian and North American *Limenitis* species are embedded in North American *Adelpha* making the genus *Adelpha* paraphyletic. We follow Willmott (2003)^15^ and the more recent updates by Prudic et al. (2008)^60^ and Rush et al. (2023)^61^ for *Adelpha* taxonomy. For identifying the opsins expressed in each species using RNA-seq, adult butterflies from seven *Adelpha* and three *Limenitis* species (*A. basiloides* (n=1), *A. californica* (n=2), *A. donysa* (n=1)*, A. ethelda,* (n=1), *A. iphiclus* (n=1)*, A. leucerioides* (n=1)*, A. pithys* (n=1), *L. lorquini* (n=1)*, L. weidemeyerii* (n=1), and *L. archippus* (n=1)) were collected from various localities in Oaxaca, Mexico and the U.S.A. from 2007-2009 ([permit numbers: SEMARNAT FAUT- 0148, CONABIO B150 and CONABIO H209] Supplementary Data 1) and used for Sanger sequencing. Adult individuals from ten additional species (*A. boreas* (n=1)*, A. cocala* (n=2)*, A. cytherea* (n=2)*, A. erotia* (n=2)*, A. heraclea* (n=2)*, A. leucophthalma* (n=2)*, A. malea* (n=1)*, A. naxia* (n=4)*, A. phylaca* (n=2) and *A. serpa celerio* (n=2)) were subsequently collected from Costa Rica from 2015-2017 using aerial insect nets, or were reared on host plants on which they were found in the field [permit numbers: R- 003-2016-OT-CONAGEBIO; R-021-2016-OT-CONAGEBIO]. *Limenitis arthemis astyanax* (n=2) were collected in Florida in 2016 (Supplementary Data 1). These wild-caught adult specimens were collected into RNALater (see below). Tissues were stored in a refrigerator whenever possible in the field and stored at -20° C upon arriving at the laboratory. Specimen sex and collection locality data are given in Supplementary Data 1.

*Adelpha fessonia* pupae (n=5) were obtained from the Costa Rica Entomological Supply in October 2017 and these specimens were used for quantification of opsin expression levels using RNA-seq. The live *A. fessonia* pupae (4 males and 1 female) were allowed to eclose in the lab under the illumination of a large window as well as overhead fluorescent lights. Adult butterfly heads (eye+brain) were then dissected into RNAlater (Invitrogen) within a week of eclosion between 10 a.m. and 4 p.m. by removing and bisecting the head between the eyes, removing the walking legs, cutting the thorax in half between the middle and hind legs, and removing the abdomen then submerging the tissues in RNAlater solution.

RNA was extracted from all specimens using the Trizol (Invitrogen) extraction method. Opsin sequences for *A. iphiclus, A. leuceroides, A. basioloides, A. ethelda, A. donysa, A. pithys, L. weidemeyerii, L. lorquini, L. archippus,* and *A. californica* were obtained using RT-PCR with degenerate PCR primers and direct Sanger sequencing of the PCR products^30^. For the other samples, TruSeq RNA Sample Preparation Kit v2 (Illumina) was used to prepare libraries for 100 bp paired-end RNA-seq. Sequencing was done on a HiSeq 2500 at the UCI Genomics High-Throughput Facility.

### *De novo* transcriptome assembly

We built head tissue *de novo* Trinity assemblies for each of 22 *Adelpha* and 2 *Limenitis* species (Supplementary Data 1 and 2). Raw reads were trimmed for quality and parsed using custom scripts. To create the *de novo* transcriptome assemblies, we used the Trinity pipeline (trinityrnaseq_r2012-06-08v2) on the University of California’s Legacy High Performance Computing cluster (HPC2)^62,63^. We deposited the resulting Trinity assemblies for *A. boreas* (n=1) *, A. cocala* (n=2)*, A. cytherea* (n=2)*, A. erotia* (n=2)*, A. fessonia* (n=2)*, A. heraclea* (n=2)*, A. leucophthalma* (n=2)*, A. malea* (n=1)*, A. naxia* (n=4)*, A. phylaca* (n=2)*, A. serpa celerio* (n=2) and *Limenitis arthemis* (n=2) in Dryad (DOI: 10.5061/dryad.xd2547dr5)^64^ and the corresponding raw RNA-seq data in NCBI SRA under BioProject Accession number PRJNA1152103.

The transcriptomes were searched for opsin sequences using BLASTX and *Heliconius melpomene* opsin query sequences^53^. Sequences with identity of more than 20% and an e-value greater than 1E-10 were extracted and translated into amino acid sequence using Orf Predictor with the blastx option before testing their homology by reciprocal blast hit^65^. UV (n=34), B (n=34) and LW (n=34) opsin sequences from *Adelpha* and *Limenitis* species either obtained through Sanger or RNA sequencing were annotated and deposited in GenBank with accession numbers: OR914986-OR915048, PP958853-PP958878, PP992301-PP992313 and PQ301474 (Supplementary Data 1).

### Phylogenetic Analysis

The evolutionary history of each opsin gene family member was inferred by translating and then aligning the complete nucleotide coding sequence for each opsin. After manual inspection, the maximum likelihood method based on the HKY85 model as implemented in PhyML^66^ was used to build trees from 34 full-length coding sequences each of UV and blue opsin transcripts and 35 full-length coding sequences of LW opsin transcripts. Support for each node was evaluated out of 500 bootstrap replicates.

### Quantifying opsin gene expression in *Adelpha fessonia*

To determine the involvement of opsins in vision, we looked at the mRNA expression levels of the three *A. fessonia* visual opsins (*UVRh*, *BRh* and *LWRh*) along with expression levels of mRNAs encoding four other members of the opsin gene family (unclassified opsin, RGR-like opsin, pteropsin, and RH7)^53^ in eye + brain tissue using kallisto (v0.46.2) RNA quantification^67^. A custom BLAST database was made from the *de novo* Trinity assembly for *A. fessonia*. Protein query sequences for each opsin gene of interest were aligned against the custom BLAST database, and corrected *A. fessonia* sequences were added to the *de novo* transcriptome assembly. Transcriptome-wide gene expression in transcripts per million (TPMs) was then quantified in kallisto by matching reads from *A. fessonia* RNA-seq libraries to k-mer type in pseudoalignments as opposed to direct read alignments^67^. Expression levels were visualized by graphing the log base 2 of the TPM for each gene.

### Testing for selection in the blue opsins of *Adelpha*

*Adelpha* blue opsins have either a tyrosine or a phenylalanine at amino acid position 195. This amino acid substitution causes a spectral tuning shift in *Pieris rapae* and *Limenitis arthemis* blue-absorbing rhodopsins^40,49^. The Y195F substitution red shifts the blue opsin’s peak from ∼431 nm to ∼435 nm. To identify possible positively selected sites, we first generated opsin amino acid sequence alignments in MEGA-X^68,69^. Then we used a site-specific maximum likelihood model to test for selection. We made the assumption that the ratio of nonsynonymous to synonymous substitution rates (*ω*) is constant across the branches of the phylogeny but possibly different among sites. The multiple sequence alignments and phylogenetic trees were analyzed in EasyCodeML, a graphical frontend for PAML 4.9j (Phylogenetic Analysis by Maximum Likelihood)^70,71^. Site models used were M1a vs M2a (neutral vs positive) where support for positive selection is given if M2a provides a better fit than M1a using likelihood-ratio tests^72,73^.

### Epi-microspectrophotometry

*In vivo* reflectance spectra of butterfly eyeshine can be used to determine the absorbance spectrum of a long wavelength-absorbing visual pigment (LW rhodopsin) due to the rapid decay of the metarhodopsin photoproduct and relatively slow recovery of rhodopsin^30^. Repeated flashes of bright long-wavelength light convert a substantial fraction of LW rhodopsin to metarhodopsin, which then decays exponentially leaving the eye in a partially bleached state. With sufficient additional time in the dark there is substantial recovery of rhodopsin, so that difference spectra compared to the partially bleached state is an estimate of the absorbance spectrum of LW rhodopsin.

Spectral measurements of eyeshine from *Adelpha* are challenging because their corneas exhibit structural, iridescent reflections that are a source of substantial glare, perturbing measurements from the fully dark-adapted eye. To reduce the influence of corneal glare from *A. fessonia* (n=1 butterfly), the eye was treated for ten hours with 1 sec 566 nm flashes every 90 sec, which created a total bleach of LW rhodopsin and very bright eyeshine. Allowing rhodopsin to recover in the dark for two hours created a difference spectrum that was fit very well by a 530 nm rhodopsin template.

A similar strategy was employed for *A. californica* (n=1 butterfly). A bright partial bleach was created with forty minutes of 5 sec Schott OG590 flashes every 60 sec. A reference spectrum was measured after waiting for metarhodopsin to decay. Then the eye was treated with a single 3 sec Hoya O58 flash, and a second spectrum measured after metarhodopsin decayed. The difference spectrum was well fit by the R530 template^74^.

### Optophysiology

Photoreceptor cells of butterfly eyes exhibit intracellular pupillary responses to bright light that cause easily measured decreases in eyeshine reflectance. Spectral sensitivity functions of this pupillary response can be measured in a double-beam epi- MSP. The monitoring beam is set up to measure continuously the reflectance of eyeshine from the deep pseudopupil with long-wavelength light that itself is so far below threshold that there is no decrease in reflectance after that beam is first turned on. The stimulating beam, from a wavelength-adjustable and intensity-adjustable monochromatic source, presents flashes to the eye that are sufficiently bright to cause decreased monitoring reflectance. To eliminate stimulus artifacts in these measurements, the quantum-counting photomultiplier is covered by a cutoff filter that transmits the monitoring beam but blocks the stimulating flashes.

It is important to choose the wavelength content of the monitoring beam, so it is reflected well by the tapetum. For most nymphalids, the tapetum reflects well from the UV out to a long-wavelength cutoff ranging from 650 nm to 700 nm or more, depending on the species and the eye-region. For *A. fessonia* and *A. californica*, however, the cutoff wavelength is only 600 nm, beyond which the reflectance drops abruptly. For our experiment with an *A. fessonia* female (n=1), therefore, we chose a monitoring beam created by a 600 nm interference filter covered by a density 2.0 neutral filter. The eye region selected was the same as for the epi-MSP bleaching experiment. The stimulus-blocking filter covering the PMT was Schott OG590. Stimulating flashes of either 60 sec or 10 sec duration were delivered every 4 min, as required to achieve stable response by the end of the flash. The criterion response chosen was a 7% decrease in reflectance monitored and recorded by a digital oscilloscope. For wavelengths ranging from 340 nm to 580 nm every 10 nm, the criterion response was achieved. At the end of the experimental session, all quantum fluxes that created criterion responses were measured with a factory-calibrated Hamamatsu S1226BK photodiode. The spectral sensitivity data were computed as the reciprocal of those quantum fluxes vs. wavelength. The good fit of a 355 nm rhodopsin template to the pupillary sensitivity data shows that only the UV-sensitive 355 nm photoreceptors contribute to responses in the range 340 nm – 380 nm. Similarly, the fit of an 530 nm rhodopsin template from 490 nm – 580 nm shows that only the green-sensitive 530 nm photoreceptors contribute to long-wavelength sensitivity, confirming the MSP estimate that the LW rhodopsin of *A. fessonia* has a peak absorbance (λ_max_) of 530 nm. In the range 390 nm – 480 nm, the elevated sensitivity of data argues for a contribution from blue-sensitive rhodopsin, but its spectral position was not determined in these experiments.

### HEK293 cell culture expression, reconstitution, and UV-visible spectroscopy of BRh visual pigment

In a previous site-directed mutagenesis study of *Adelpha/Limenitis* BRh opsins, we identified a Y195F substitution which led to a ∼4 nm spectral red shift in peak absorbance of the reconstituted photopigment relative to the wildtype BRh opsin that had a Y at site 195^49^. In that study, however, we did not report data from a wild-type BRh opsin sequence with F at site 195. Since *A. fessonia* and *A. basiloides* differ at this site, we decided to report below the outcome of expressing a wildtype BRh opsin which is very similar in sequence to *A. fessonia* and which has a F amino acid at site 195, namely that of *L. lorquini* (with 321 identical amino acids out of 330). Total RNAs were extracted from a single adult *L. lorquini* head using Trizol (Invitrogen). cDNAs were then synthesized using a Marathon cDNA amplification kit (BD Biosciences, Franklin Lakes, NJ). The *BRh* coding region was cloned into pGEM-T easy vector (Promega). A short oligonucleotide sequence encoding the 1D4 epitope of bovine rhodopsin (STTVSKTETSQVAPA) was added before the stop codon. The tagged cDNA fragments were subcloned into expression vector pcDNA3.1(+) (Invitrogen). Transient transfection of HEK293 cells by plasmid DNA was carried out by Lipofectamine 2000 (Invitrogen). The cells were replated 2 days after transfection and treated with 1 mg/mL G418 sulfate (EMD Chemicals Inc., Gibbstown, NJ) for 2 weeks. Twelve cell clones were chosen, expanded, and screened by western blotting. The clone having the highest expression level was then expanded to 15 plates of cells. The cultured cells were incubated with 1 mM 11*-cis*-retinal for 2 days and collected by centrifugation under dim red light. The collected cells were incubated with 40 mM 11*-cis*-retinal for 1 h at 4 °C. The reconstituted visual pigments were extracted using 1% n-dodecyl b–D-maltoside (DDM) (Sigma-Aldrich, Saint Louis, MO) in 10 ml of extraction buffer (pH 6.7) (250 mM sucrose, 120 mM KCl, 10 mM MOPS, 5 mM MgCl2, 1 mM DTT, 1 Roche protease inhibitor cocktail tablet) (Vought et al. 2000) via gentle rotation for 1 h at 4 °C. The visual pigments were then purified by immunoaffinity chromatography. Briefly, the crude extract was mixed with Sepharose beads conjugated with 1D4 IgG (University of British Columbia, Canada) overnight at 4 °C. The beads were then washed with 50 ml of wash buffer, pH 6.6 (20% glycerol, 120mM KCl, 5mM MgCl2, 50mM HEPES, and 0.1% DDM) (Vought et al. 2000). The visual pigments were eluted with 50 mM competing peptide (Quality Controlled Biochemicals, Hopkinton, MA) in wash buffer and measured in a Hitachi U-3310 UV-Vis spectrophotometer at 0°C. Spectral data represent the average of 5–8 scans. The expression experiment was repeated three times for the *L. lorquini* blue opsin, twice for the Y195F construct and four times for the *L. arthemis* wildtype blue opsin. The purified photopigment experiment with the highest optical density was selected for further analysis. λ_max_ was estimated by least-squares fitting of the data to a visual pigment template^75^.

### Reflectance of Adelpha fessonia and A. basiloides wings

Wings of *A. fessonia* and *A. basiloides* were obtained as dry specimens from the collection of the Museo de Zoología Alfonso L. Herrera in the Universidad Nacional Autónoma de México. Dry samples were rehydrated overnight before wings were excised. Wing reflectance measurements of 8 males and 8 females each of *A. fessonia* and 8 females and 11 males each of *A. basiloides* were taken (Supplementary Data 3).

Briefly, after inspecting individual wings for damage, measurements were taken of intact dorsal orange spots and white stripes as these color patches are likely to elicit responses from conspecifics or avian predators. The fixed probe holder (Ocean Optics RPH-1) was placed so that the axis of the illuminating and detecting bifurcating fiber (Ocean Optics 400-7-UV/VIS) was 45 degrees to the plane of the wings. Illumination was provided by a deuterium-halogen tungsten lamp (Ocean Optics DH-2000) with the reflectance spectra measured by an Ocean Optics USB2000 spectrometer. Measurements were recorded on Ocean View software v. 1.6.7 and smoothed and plotted using the r package, “pavo” and r version 4.3.3 (citation)^76,77^. Specimen ID, sex and collection locality information for *A. fessonia* and *A. basiloides* butterflies used in wing reflectance measurements are given in Supplementary Data 3.

### Visual modeling of *Adelpha* wings through the eyes of predator and prey

We used the receptor-noise model to predict whether *Adelpha fessonia* and avian predators can distinguish *A. fessonia* from *A. basiloides* orange patches or *A. fessonia* from *A. basiloides* white stripes^78^. The reflective distribution of the measured stimuli from the reflectance measurements were plotted on a triangular *A. fessonia* color space and on the tetrahedral color spaces of ultraviolet- and violet-sensitive birds. Using the “vismodel” function from “pavo”, we calculated the quantum catches at each photoreceptor. For the avian models, we used normalized rhodopsin templates for chicken (418 nm, 453 nm, 507 nm, and 571 nm) and blue tit (372 nm, 449 nm, 502 nm, and 563 nm) visual systems^79^ for chromatic distances. We used the normalized double-cone sensitivity of the blue tit (563 nm) and chicken (571 nm) for luminance distances of the UV-sensitive and violet-sensitive avian visual systems, respectively. For *A. fessonia*, we used λ_max_ values corresponding to 355 nm, 431 nm, and 530 nm, respectively, for the chromatic distances and the 530 nm photoreceptor for luminance distances. The color spaces were generated using the “plot” and “colspace” functions^76^.

Chromatic and achromatic distances between the orange patches or the white stripes of *A. fessonia* and *A. basiloides* were calculated using the “bootcoldist” function. Bootcoldist allowed us to perform a 1000 bootstrap replicates of contrast (JND) with a 95% confidence interval to determine if the color distance measurement exceeds a theoretical discrimination threshold^76^. A JND threshold of one is typically used for laboratory experiments. However, spectra separated by a JND threshold of one could still be impossible to distinguish between each other especially under field conditions^80^.

We use a relative receptor density of 1:2:2:4 (ultraviolet:short:medium:long-wave)^78,81^ along with a Weber fraction of 0.1 for chromatic^82–84^ and 0.18 for achromatic receptors for the UV-sensitive bird model^85^. For the violet-sensitive bird model, we use a relative receptor density of 1:2:4:4 (short:short:medium:long)^86^ as well as a Weber fraction of 0.06 for both chromatic^87,88^ and 0.22 for achromatic receptors^85,89^.

Over a period of years, we tried to obtain additional pupae of *A. fessonia* from our Costa Rican source to perform *in situ* hybridization on this species to quantify relative receptor densities but were unsuccessful. Therefore, *Adelpha fessonia*’s relative receptor density of 0.15, 0.13, 1 (UV:short:long) was approximated from a *Limenitis arthemis* opsin *in situ* hybridization image^49^. Photoreceptor density was manually performed by counting a random sample of 102 intact ommatidia and normalizing the photoreceptors in proportion to the long-wave sensitive photoreceptor. Weber fraction estimates based on physiological or behavioral experiments do not exist for butterflies. Therefore, we used weber fractions derived from honeybee electrophysiological and behavioral experiments as follows: 0.13 (UV), 0.06 (B), 0.12 (LW) for chromatic^90,91^ and 0.16 for achromatic receptors^92^. Parameters for each of the visual models, which are based on anatomical, physiological or behavioral experiments, are given in Table 1. Illumination for all visual models was set to D65 and the average reflectance spectra of *Adelpha* brown color measurements (n=29 independent measurements) was used for the background. Calculated color distances were then plotted using rstudio’s “plot” function.

### Statistics and Reproducibility

Samples sizes for individual measurement are indicated in each figure legend, within the figure itself, or in Supplementary Data 1 and 3. Replicates are biological replicates. Cell culture experiments were performed 2-4 times (biological replicates) with the experiment yielding the highest optical density of the blue-absorbing rhodopsin selected for λ_max_ estimation. Each panel in Fig. 5 represents the average of 5-7 scans (technical replicates) from the individual cell culture experiment with the highest yield. Bootstrapping statistics for the phylogenetic analyses were calculated using default settings in Geneious Prime v. 2024.0.5 (https://www.geneious.com)^93^ and 95% bootstrapped confidence intervals for the JNDs were calculated using default settings in pavo 2^76^.

## Data availability

The opsin sequence data are available in Genbank under accession numbers OR914986-OR915048, PP958853-PP958878, PP992301-PP992313 and PQ301474, the transcriptome assemblies, transcriptome fasta files and wing reflectance spectra, along with raw data used to make figures, are deposited in Dryad (DOI: 10.5061/dryad.xd2547dr5)^64^ and the raw RNA-seq reads are available in NCBI SRA under BioProject Accession number PRJNA1152103.

## Code Availability

Geneious Prime v. 2024.0.5 software is commercially-available at https://www.geneious.com. kallisto^67^ is available at http://pachterlab.github.io/kallisto/. pavo 2^76^ is available at https://cran.r-project.org/web/packages/pavo/index.html. r version 4.3.3 is available at https://www.r-project.org/^94^ .

## Author contributions

A.D., G.D.B, S.P.M, and A.D.B. designed research; A.D., G.D.B., A.M.M., F.Y., R.I.H, J.P.L, A.G.R.O., A.L.-M., J.L.B. performed research; A.D., G.D.B., F.Y., A.M.M., A.G.R.O., and A.D.B. analyzed data; A.D. and A.D.B. wrote the paper.

## Acknowledgments

We thank Omar Ávalos, Francesca Frentiu, Fred Gagnon, Luis Gonzáles Cota, Blanca Claudia Hernández, Maita Kuvhenguhwa, Máximo Martínez, Gordon Pratt, Marysol Trujano, and Andrew Warren for assistance with specimen collection, the Costa Rica Entomological Supply for providing pupae, and Maita Kuvhenguhwa and Zachary Johnston for technical support. Thanks also to Daniel Osorio, Kentaro Arikawa, Natalie Hempel de Ibarra, Gregor Belušič and two anonymous reviewers for feedback on the visual models, and Sarah Hoenicke Flores for proofreading the manuscript. This work was supported by NSF grants DEB-1342706 to R.I.H. and DEB-1342759 and IOS- 1656260 to A.D.B. and a HFSP grant RGP005/2023 (https://doi.org/10.52044/HFSP.RGP0052023.pc.gr.168591)^95^ to A.D.B.

## Supplementary information

**Supplementary Figure 1.** Reflectance spectra of orange and white dorsal wing colors of male and female *Adelpha fessonia* and *A. basiloides.* The number of individual butterflies measured for each comparison are indicated as numbers in parentheses with *A. fessonia* on the left and *A. basiloides* on the right.

**Supplementary Data 1.** Specimen collecting locality, sex, sequencing method, and opsin Genbank accession numbers.

**Supplementary Data 2.** RNA-seq library assembly stats.

**Supplementary Data 3.** Specimen collecting locality and sex for wing reflectance measurements.

