## Supplementary Figure 1 for "Mimetic butterfly wings through mimetic butterfly eyes"

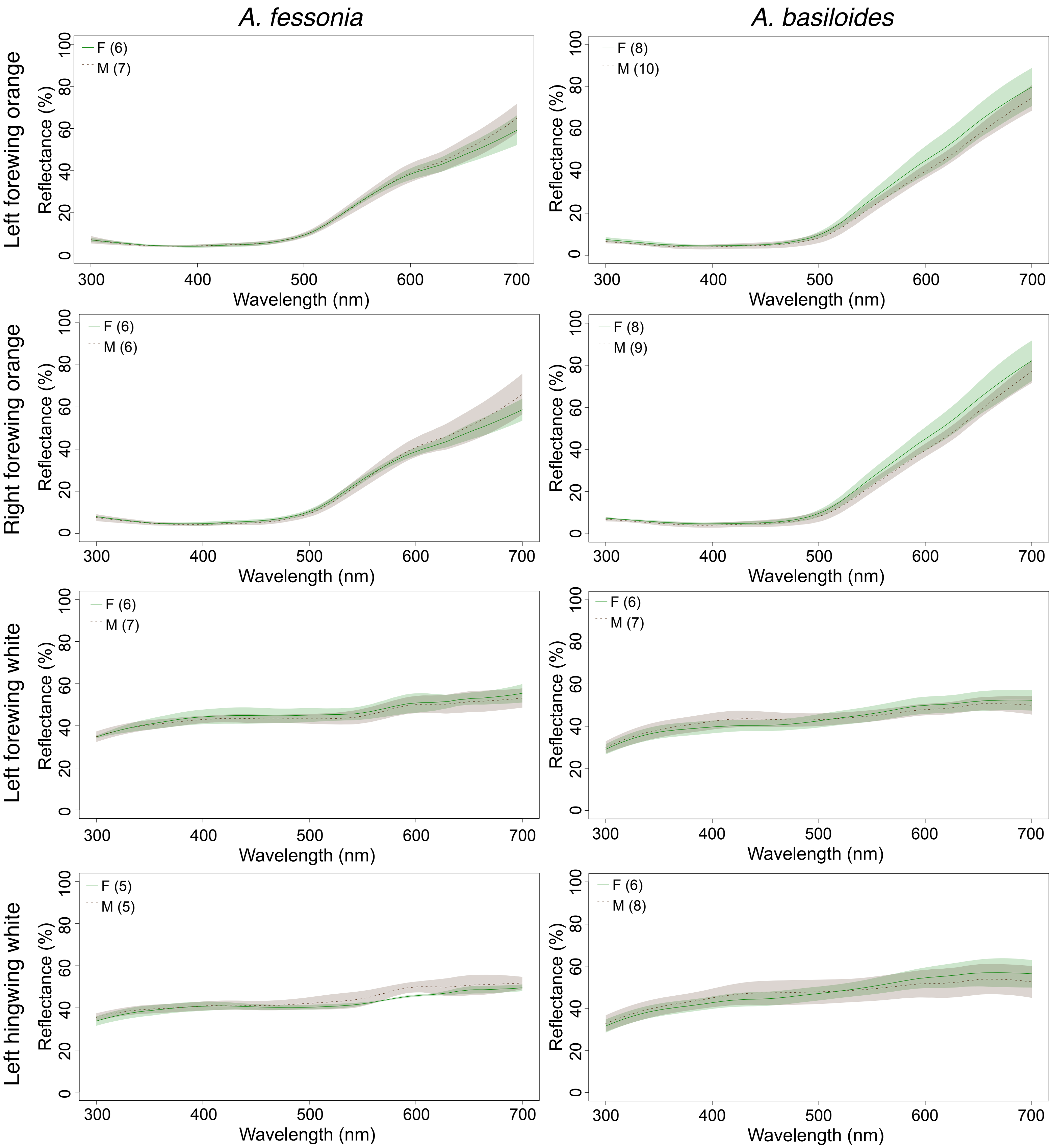


**Supplementary Figure 1.** Reflectance spectra of orange and white dorsal wing colors of male and female *Adelpha fessonia* and *A. basiloides*. The number of individual butterflies measured for each comparison are indicated as numbers in parentheses with *A. fessonia* on the left and *A. basiloides* on the right.
